# Can sleep protect memories from catastrophic forgetting?

**DOI:** 10.1101/569038

**Authors:** Oscar C. González, Yury Sokolov, Giri P. Krishnan, Maxim Bazhenov

## Abstract

Continual learning remains to be an unsolved problem in artificial neural networks. Biological systems have evolved mechanisms by which they can prevent catastrophic forgetting of old knowledge during new training and allow lifelong learning. Building upon data suggesting the importance of sleep in learning and memory, here we test a hypothesis that sleep protects memories from catastrophic forgetting. We found that training in a thalamocortical network model of a “new” memory that interferes with previously stored “old” memory may result in degradation and forgetting of the old memory trace. Simulating NREM sleep immediately after new learning leads to replay, which reverses the damage and ultimately enhances both old and new memory traces. Surprisingly, we found that sleep replay goes beyond recovering old memory traces that were damaged by new learning. When a new memory competes for the neuronal/synaptic resources previously allocated to the old memory, sleep replay changes the synaptic footprint of the old memory trace to allow for the overlapping populations of neurons to store multiple memories. Different neurons become preferentially supporting different memory traces to allow successful recall. We compared synaptic weight dynamics during sleep replay with that during interleaved training – a common approach to overcome catastrophic forgetting in artificial networks – and found that interleaved training promotes synaptic competition and weakening of reciprocal synapses, effectively reducing an ensemble of neurons contributing to memory recall. This leads to suboptimal recall performance compared to that after sleep. Together, our results suggest that sleep provides a powerful mechanism to achieve continual learning by combining consolidation of new memory traces with reconsolidation of old memory traces to minimize memory interference.

## Introduction

Animals and humans are capable of continuous, sequential learning. In contrast, modern artificial neural networks suffer from the inability to perform continual learning (French, 1999; Hassabis et al., 2017; Hasselmo, 2017; Kirkpatrick et al., 2017; Ratcliff, 1990). Training new task results in interference and catastrophic forgetting of old memories (French, 1999; Hasselmo, 2017; McClelland et al., 1995; Ratcliff, 1990). Several attempts have been made to overcome this problem including (1) explicit retraining of all previously learned memories – interleaved training (Hasselmo, 2017), (2) using generative models to reactivate previous inputs (Kemker and Kanan, 2017), (3) artificially “freezing” subsets of synapses critical for the old memories (Kirkpatrick et al., 2017). These solutions help prevent new memories from interfering with the previously stored old memories, however they either require explicit retraining of all old memories using the original data or have limitations on the types of trainable new memories and network architectures (Kemker et al., 2017). How biological systems avoid catastrophic forgetting and reduce interference of the old and new memories supporting continuous learning remains to be understood. In this paper, we propose a mechanism for how sleep modifies the synaptic connectivity matrix to minimize interference of competing memory traces enabling continual learning.

Sleep has been suggested to play an important role in memory and learning (Oudiette and Paller, 2013; Paller and Voss, 2004; Rasch and Born, 2013; Stickgold, 2013; Walker and Stickgold, 2004; Wei et al., 2016; Wei et al., 2018). Specifically, the role of stage 2 (N2) and stage 3 (N3) of Non-Rapid Eye Movement (NREM) sleep has been shown to help with the consolidation of newly encoded memories (Paller and Voss, 2004; Rasch and Born, 2013; Stickgold, 2013; Walker and Stickgold, 2004). The mechanism by which memory consolidation is influenced by sleep is still debated, however, a number of hypotheses have been put forward. Sleep may enable memory consolidation through repeated reactivation or replay of specific memory traces during characteristic sleep rhythms such as spindles and slow oscillations (Clemens et al., 2005; Ladenbauer et al., 2017; Marshall et al., 2006a; Oudiette et al., 2013; Paller and Voss, 2004; Rasch and Born, 2013; Wei et al., 2016; Wei et al., 2018; Xu et al., 2019). Memory replay during NREM sleep could help strengthen previously stored memories, map memory traces between brain structures and prevent interference. Previous work using electrical (Ladenbauer et al., 2017; Marshall et al., 2006a; Marshall et al., 2004) or auditory (Ngo et al., 2013) stimulation showed that increasing neocortical sleep oscillations during NREM sleep resulted in improved consolidation of declarative memories. Similarly, spatial memory consolidation has been shown to improve following cued reactivation of memory traces during slow-wave NREM sleep (Oudiette et al., 2013; Oudiette and Paller, 2013; Paller and Voss, 2004; Papalambros et al., 2017). Our recent computational studies found that slow-wave sleep dynamics can lead to replay and strengthening of the recently learned memory traces (Wei et al., 2016; Wei et al., 2018). These studies point to the critical role of sleep in memory consolidation.

Can neuroscience inspired ideas help solve the catastrophic forgetting problem in artificial neuronal networks? The most common machine learning training algorithm – backpropagation (Kriegeskorte, 2015; Rumelhart et al., 1986) – is very different from plasticity rules utilized by the brain networks. Nevertheless, we have recently seen a number of successful attempts to implement high level principles of biological learning in artificial network designs, including implementation of the ideas from “Complementary Learning System Theory” (McClelland et al., 1995), according to which the hippocampus is responsible for the fast acquisition of new information, while the neocortex would more gradually learn a generalized and distributed representation. These ideas led to interesting attempts of solving catastrophic forgetting problem in artificial neural networks (Kemker and Kanan, 2017). While few attempts have been made to implement sleep in artificial networks, one study suggested that sleep-like activity can increase storage capacity in artificial networks (Fachechi et al., 2019). We recently found that implementation of the sleep-like phase in the artificial networks trained using backpropagation can dramatically reduce catastrophic forgetting, as well as improve generalization performance and transfer of knowledge (Krishnan, 2019).

In this new study, we used a biophysically realistic thalamocortical network model to identify the network level mechanisms of how sleep promotes consolidation of newly encoded memory traces and reverses damage to older memories. We considered a scenario of procedural memories which are directly encoded in the populations of neocortical neurons. Our model predicts that a period of slow-wave sleep, following training of a new memory in awake, can promote replay of both old and new memory traces thereby preventing forgetting and increasing recall performance. We show that sleep replay results in the fine tuning of the synaptic connectivity matrix representing the interfering memory traces to allow for the overlapping populations of neurons to store multiple competing memories.

## Results

The thalamocortical network model included two main structures (cortex and thalamus), each comprised of excitatory and inhibitory cell populations (figure 1A). Both populations included well characterized excitatory-inhibitory circuits. In the thalamus, excitatory thalamocortical neurons (TCs) received excitatory connections from cortical excitatory neurons (PYs), and inhibitory connections from thalamic reticular neurons (REs). RE neurons were interconnected by inhibitory synapses and received excitatory connections from PYs and TCs. PYs in the cortex received excitatory input from thalamus as well as excitatory inputs from other PYs and inhibitory inputs from cortical inhibitory interneurons (INs). INs received excitatory inputs from PYs in addition to the excitatory connections from TCs. All PYs had a 60% probability of connecting to neighboring PYs (PY-PY radius was set to 20 neurons). Figure 1D (left) shows the adjacency matrix for the cortical PYs that arise from these two restrictions. The initial strength of the synaptic connections between PYs was Gaussian distributed (figure 1D right). The model simulated transitions between awake and sleep (figure 1B/C) by modeling effects of neuromodulators (Krishnan et al., 2016). PY-PY connections were plastic and regulated by spike-timing dependent plasticity (STDP) that was biased for potentiation during awake state to simulate the higher level of acetylcholine (Blokland, 1995; Shinoe et al., 2005; Sugisaki et al., 2016).

**Figure 1.**
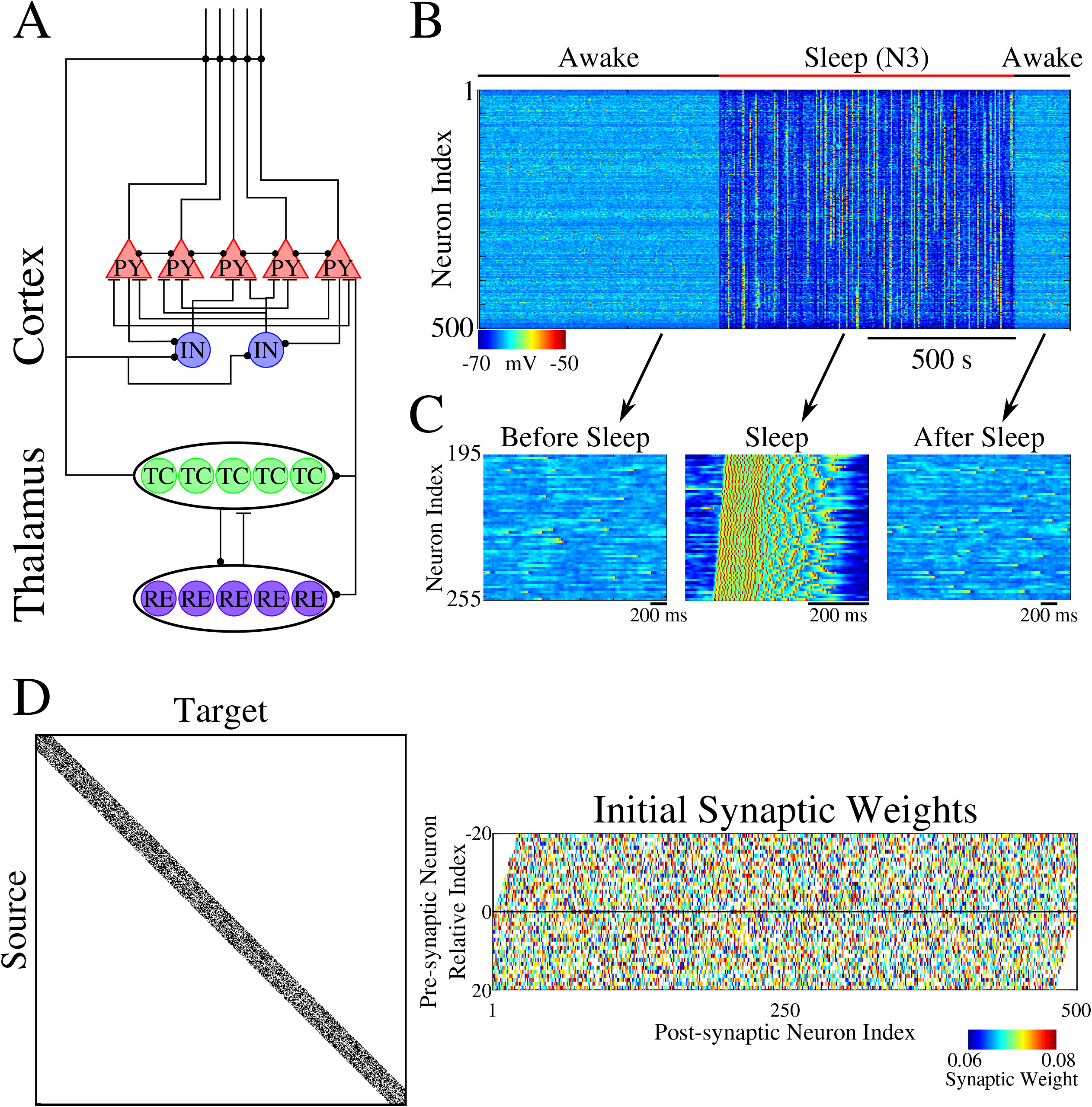
Network architecture and baseline behavior. **A**, Network schematic shows the basic network architecture (PY: excitatory pyramidal neurons; IN: inhibitory interneurons; TC: excitatory thalamocortical neurons; RE: inhibitory reticular neurons). Excitatory synapses are represented by lines terminating in a dot, while inhibitory synapses are represented by lines terminating in horizontal bars. **B**, Behavior of a control network exhibiting wake-sleep transitions. Color represents the voltage of a neuron at a given time during the simulation (dark blue – hyperpolarized potential; light – depolarized potential; red - spike). **C**, Zoom-in of a subset of neurons in the network in B (time indicated by arrows). Left and right panels show spontaneous activity during awake-like state before and after sleep, respectively. Middle panel shows example of slow wave like activity during sleep. **D**, Left panel shows adjacency matrix for the network in B. Black spots represent synaptic connections, while white represents a lack of synaptic connection between the source and target neuron. Right panel shows the initial synaptic weight matrix for the network in B. The X-axis indicates index of a post-synaptic neuron onto which the connection is made, while the Y-axis indicates the relative index of the presynaptic neuron. The sign of the presynaptic relative index (negative or positive) corresponds to the neurons to the left or right of the post-synaptic neuron, respectively. The color in this plot represented the strength of the AMPA connection between neurons, with white indicating lack of synaptic connections.

### Sequential training of spatially separated memory sequences does not lead to interference

We trained two memory sequences sequentially (first S1 and then S2) in spatially distinct regions of the network as shown in figure 2A. Each memory sequence was represented by the spatio-temporal pattern of 5 sequentially activated groups of 10 neurons per group. In the following we will use labels – A, B, C, D, E – assigned to each group of neurons, so each sequence could be labeled by a unique “word” of such “letters.” The first sequence (S1) was trained in the population of neurons 200-249 (figure 2B) with groups of 10 neurons that were ordered by increasing neuron numbers (A-B-C-D-E). Training S1 resulted in an increase of synaptic weights between participating neurons (figure 2D, top) and an increase in performance on sequence completion, which was defined as a probability to complete the entire sequence (word) after only the first group of neurons (first letter) was activated (figure 2B/C, top). While the strength of the synapses in the direction of S1 increased, synapses in the opposite direction showed a reduction of strength consistent with the STDP rule used in this network (see *Methods and Materials*). The second sequence (S2) was trained for an equal amount of time as S1 in the neuron population 350-399 (W-V-X-Y-Z, figure 2B bottom). Training of S2 also resulted in synaptic weight changes (figure 2D, middle) and improvement in performance (figure 2B/C, bottom). Importantly, training of S2 did not interfere with the weight changes encoding S1 because both sequences involved spatially distinct populations of neurons (compare figure 2D, top and middle).

**Figure 2.**
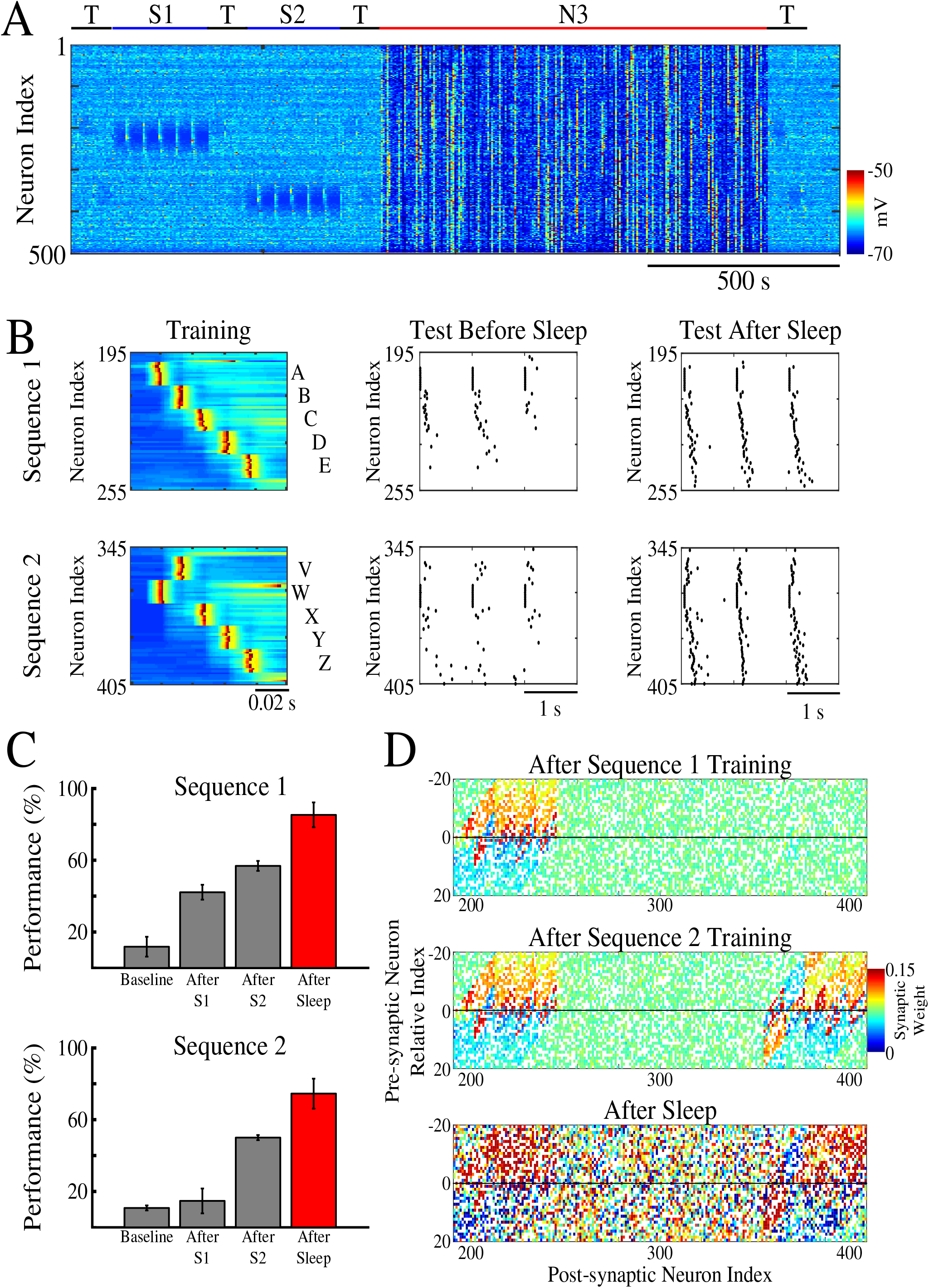
Two spatially separated memory sequences show no interference during training and both can be strengthened by subsequent sleep. **A**, Network activity shows periods of testing (T), training of two spatially separated memory sequences (S1/S2), and sleep (N3). Color indicates voltage of neurons at a given time. **B**, Left panels show an example of training of sequence 1 (S1, top) and sequence 2 (S2, bottom). Middle panels show examples of testing of both sequences prior to sleep. Right panels show examples of testing after sleep. Note, after sleep, both sequences show better sequence completion. **C**, Performance of sequences 1 and 2 before any training (baseline), after sequence 1 training (after S1), after sequence 2 training (after S2), and after sleep (red). **D**, Synaptic weight matrices show changes in synaptic weights in the regions trained for sequences 1 and 2. Top panel shows weights after training S1; middle panel shows weights after training S2; bottom one shows weights after sleep. Color indicates strength of AMPA synaptic connections.

After successful training of both memories, the network went through a period of slow-wave (N3) sleep when no stimulation was applied. After sleep, synaptic weights for both memory sequences revealed strong increases in the direction of their respective patterns and further decreases in the opposing directions (figure 2D, bottom). In line with our previous work (Wei et al., 2018), these changes were a result of memory sequence replays during Up states of slow-waves. Synaptic strengthening increased the performance on sequence completion after sleep (figure 2B, right; 2C, red bar).

### Training overlapping memory sequences results in interference

We next tested whether our network model shows interference when a new sequence (S1*) (figure 3A) is trained in the same population of neurons as the earlier old sequence (S1). S1* included the exact same population of neurons as S1, but was trained in the opposite direction to S1, that is, the stimulation group order was E-D-C-B-A (figure 3B). S2 was once again trained in a spatially distinct region of the network (figure 3A/B). Testing for sequence completion was performed immediately after each training period. This protocol can represent two somewhat different learning scenarios: (a) two competing memory traces (older S1 and newer S1*) are trained sequentially; (b) the first (old S1) memory is trained and then consolidated during sleep (as in figure 2) followed by the training of the second (new S1*) memory. We explicitly tested both scenarios and they behaved identically, so in the following we discuss the simpler case of two sequentially trained memories.

**Figure 3.**
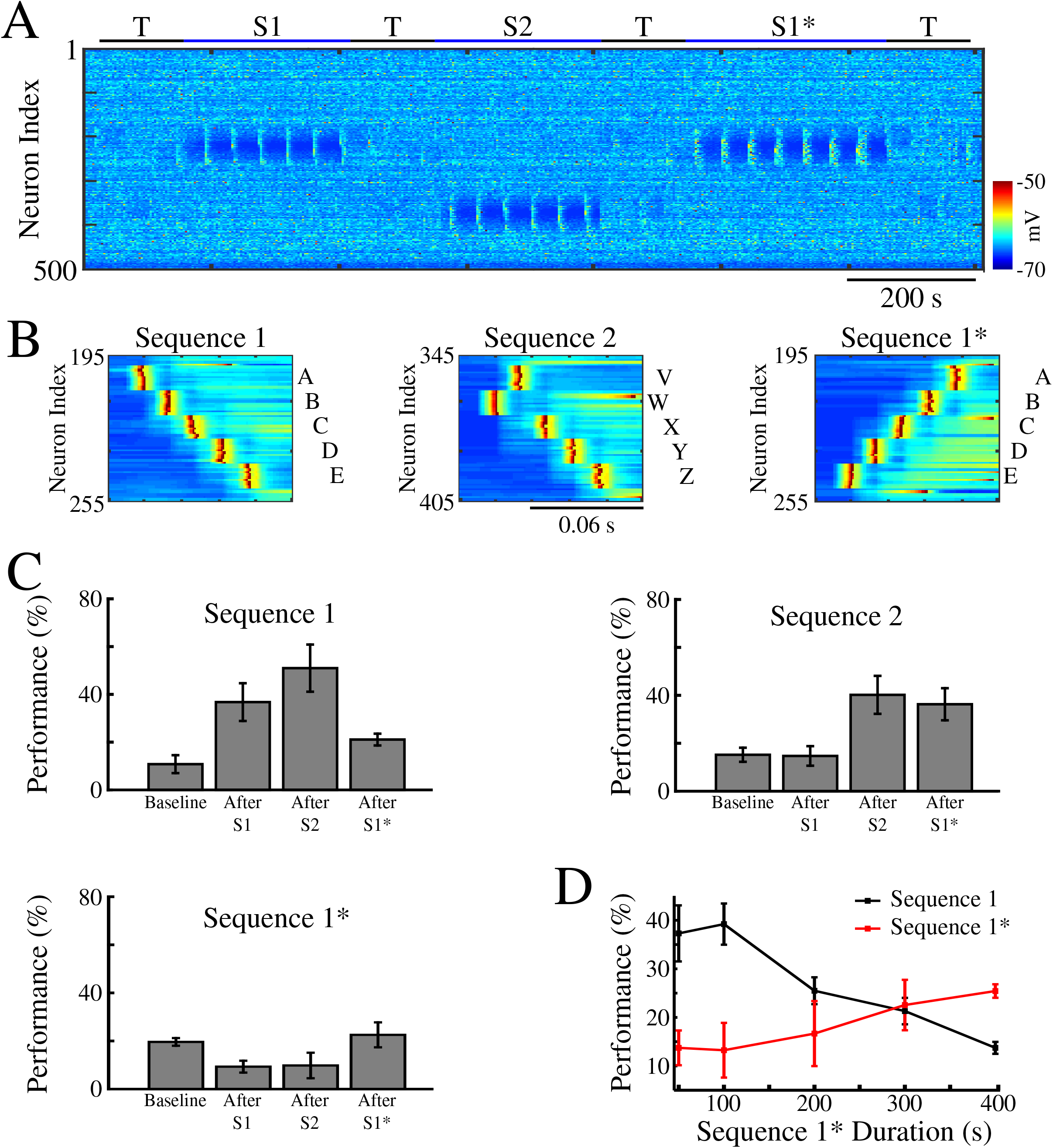
Training of overlapping memory sequences results in catastrophic interference. **A**, Network activity during training and testing periods for three memory sequences in awake-like state. Note, sequence 1 (S1) and sequence 1* (S1*) are trained over the same population of neurons. Color indicates the voltage of the neurons at a given time. **B**, Examples of sequence training protocol for S1 (left), S2 (middle), and S1* (right). **C**, Performances for the three sequences at baseline, and after S1, S2 and S3 training. Training of S1* leads to reduction of S1 performance. **D**, Performance of S1 (black) and S1* (red) as a function of S1* training duration. Note that longer S1* training increases degradation of S1 performance.

Similar to the previous results, training of S1 increased performance of S1 completion (figure 3C, top/left). It also led to decrease in performance for S1* below its baseline level in the “naive” network (figure 3C, bottom/left). (Note that even a naive network displayed some above zero probability to complete a sequence depending on the initial strength of synapses and spontaneous network activity). Training S2 led to increase in S2 performance. S1 performance also increased, probably because of the random reactivation of S1 in awake, but it was not significant. Subsequent training of S1* resulted in both a significant increase in S1* performance and a significant reduction of S1 performance (figure 3C). To evaluate the impact of S1* training on S1 performance, we varied the duration of S1* training (figure 3D). Increasing length of S1* training correlated with a reduction of S1 performance up to the point when S1 performance was reduced back to baseline level (Fig 3D, 400 s training duration of S1*). This suggests that sequential training of two memories competing for the same populations of neurons results in memory interference and catastrophic forgetting of the earlier memory sequence.

### Interleaved training can recover the old memory after damage

One well known solution to avoid catastrophic forgetting in neural networks is to use a new training dataset that includes both old and new training patterns (McClelland et al., 1995). Thus, we next examined if an interleaved presentation of S1 and S1* can reverse the damage and rescue S1. In the interleaved training protocol, S1 and S1* stimulation patterns are interleaved at subsequent trails (S1->S1*->S1->S1*->…) (figure 4 A/B).

**Figure 4.**
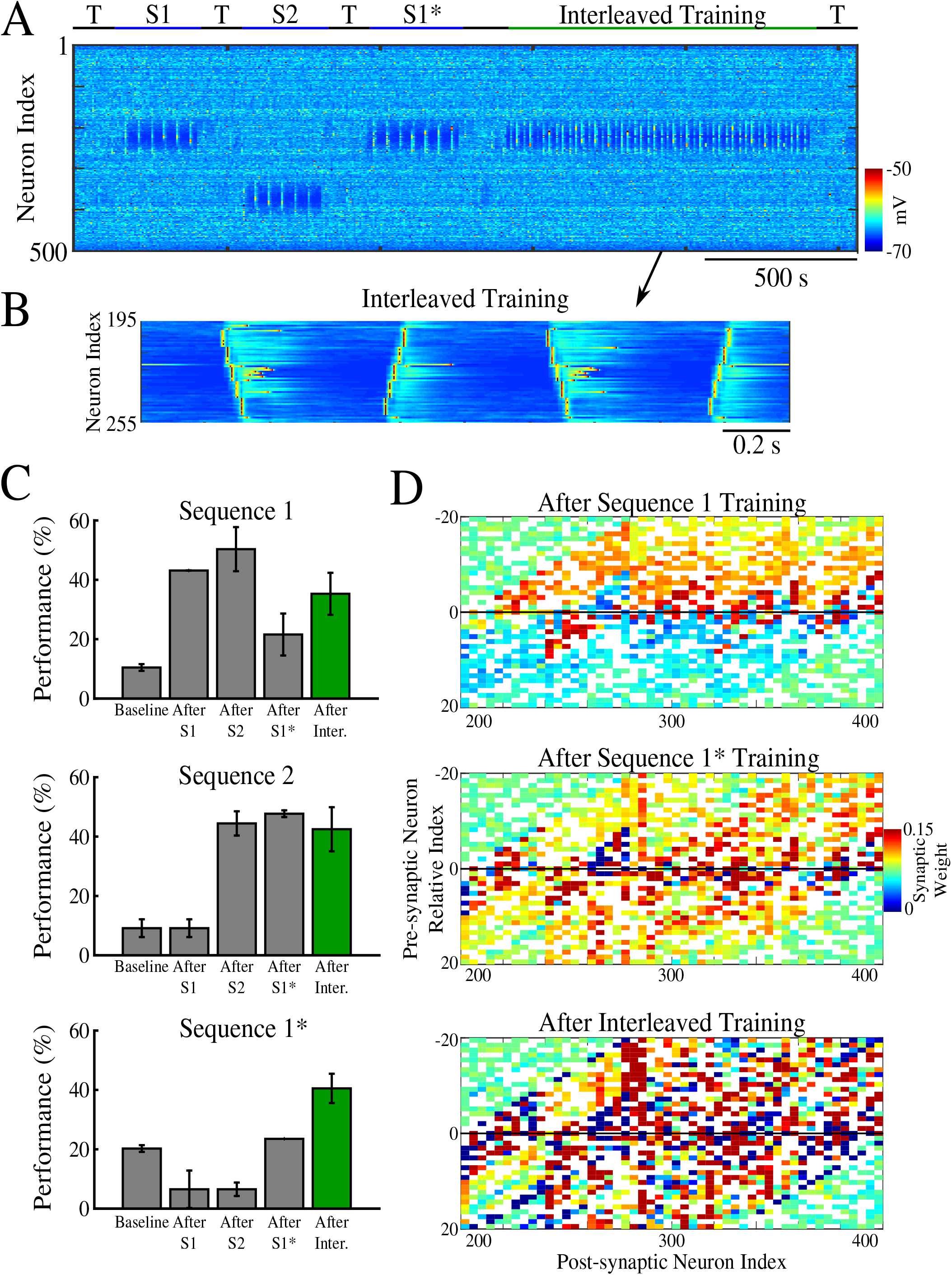
Interleaved training of the “old” and “new” memory sequences recovers the “old” sequence and improves performance for both memories. **A**, Network activity during sequential training of the memory sequences S1->S2->S1* (blue bars) followed by interleaved training of S1/S1* (green bar). **B**, Example of stimulation protocol used for interleaved training of S1/S1*. **C**, Testing of S1, S2, and S1* shows significant increase in performance of S1 and S1* after interleaved training (green). **D**, Weight matrices show changes after initial sequential training of S1 and S1* and after interleaved S1/S1* training.

First, using the same protocol as described in the figure 3, we trained the sequences sequentially (S1->S1->…->S2->S2->…->S1*->S1*->…). This, once again, resulted in initial increase in both S1 and S2 performance followed by a reduction of S1 performance after training of S1* (figure 4C). Analysis of the synaptic connectivity matrix revealed the synaptic weight dynamics behind the change in performance (figure 4D). After S1 training, synaptic weights between neurons representing S1 increased in the direction of S1 training and decreased in the opposite network direction (figure 4D, top). Training of S1* resulted in an increase of synaptic weight in the direction of S1* while reducing the strength of synapses in the direction of S1 (figure 4D, middle). After this initial sequential training phase, interleaved training was applied. Interleaved training led to further changes in the synaptic connectivity and performance increase for both S1 and S1* sequences (figure 4C, green). We observed an increase of synaptic strength in subsets of synapses representing S1 and S1* patterns (figure 4D, bottom). These results suggest that in the thalamocortical model, similar to the artificial neural networks, interleaved training of the interfering patterns can enable successful learning of the both patterns.

### Sleep enables replay and performance improvement for overlapping memories

To test if replay of the memory traces during simulated sleep can protect them from interference, we next modeled slow-wave sleep (N3) after the patterns S1/S2/S1* were trained sequentially in awake state (S1->S1->…->S2->S2->…->S1*->S1*->…) (figure 5A, red bar), as described in the previous sections (figures 2A, 4A). During the sleep phase, no stimulation was applied and the network properties were changed by increasing K^+^ leak currents and AMPA synaptic currents in all cortical neurons (ref: (Krishnan et al., 2016)) to generate spontaneous slow-waves. Figure 5B shows raster plots of the spiking activity before vs after the sleep, which revealed improvements in sequence completion after sleep. We found that sleep was not only able to reverse the damage caused to S1 following S1* training, but it was also able to enhance all previously trained memory sequences S1/S2/S1* (figure 5C, red bar). Analysis of the cumulative sum of full sequence replays throughout the course of sleep, defined as a total count of all events when a given sequence was spontaneously completed (e.g., A->B->C->D->E), revealed similar number of replays for either S1 or S1* (figure 5D, top). Importantly, for each two populations of neurons (each two letters) within a sequence, the number of single transitions in one direction (e.g., A->B) was similar to that in opposite direction (B->A), which ensured similar amounts of synaptic weight changes for both sequences during sleep (figure 5D, bottom).

**Figure 5.**
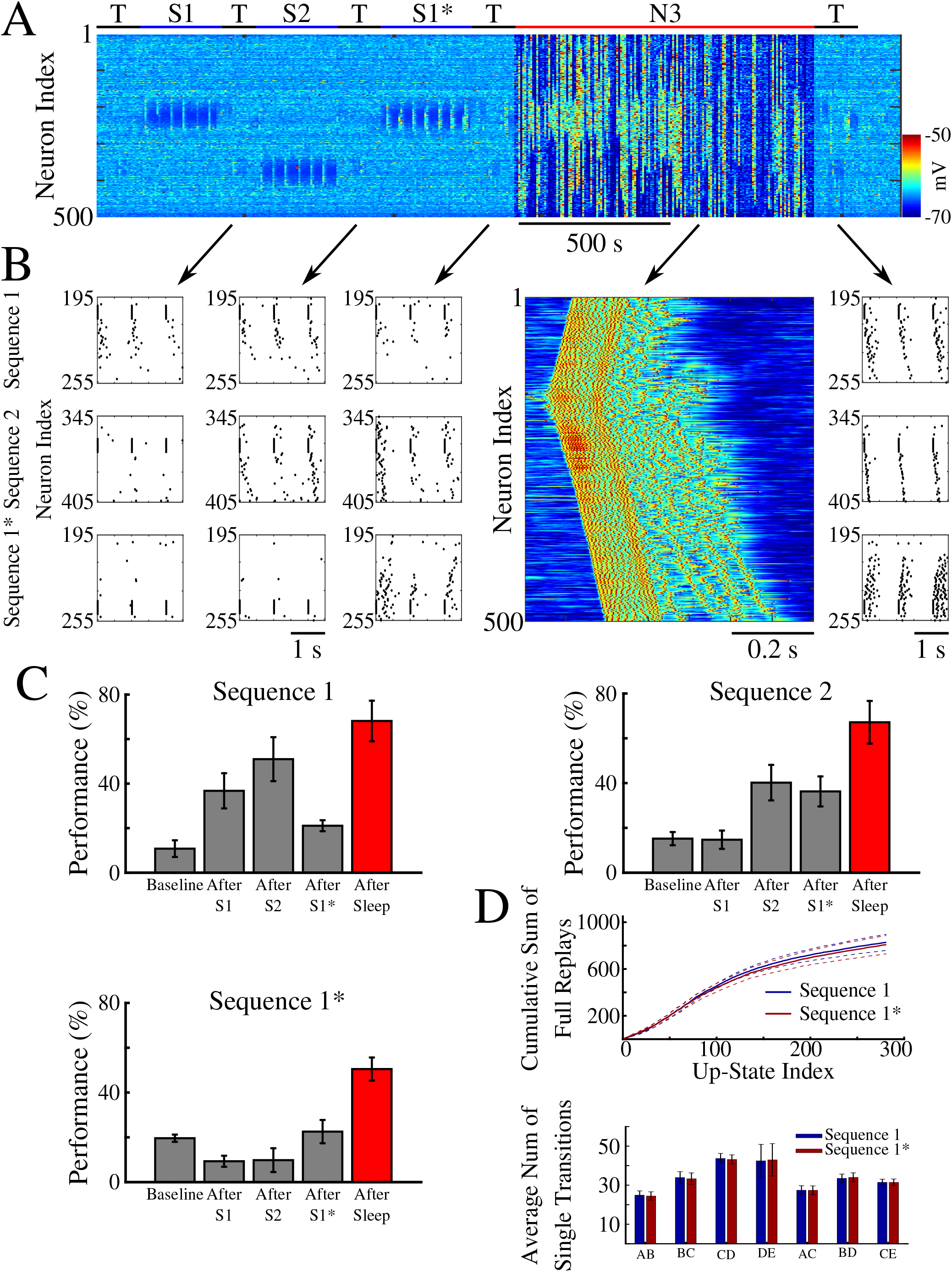
Sleep recovers the “old” memory sequence and improves performance for all three memories. **A**, Network activity during sequential training S1->S2->S1* (blue bars) followed by N3 sleep (red bar). No stimulation was applied during sleep. **B**, Examples of testing periods for each trained memory at different times. The top row corresponds to the testing of sequence 1 (S1), middle is testing of sequence 2 (S2), and bottom is testing of sequence 1* (S1*). **C**, Testing of S1, S2, and S1* shows damage to S1 after training S1*, and increase in performance of all three sequences after sleep (red bar). **D**, Top, Cumulative sum of the total number of full sequence replays during sleep. Solid lines indicate means and broken lines are SEM; Bottom, Average number of single transition replays between adjacent population of neurons (letters) over the total duration of sleep for both sequences.

### Synaptic weights dynamics during sleep replay vs interleaved training

In order to understand how sleep replay affects S1* and S1 memory traces to allow enhancement of both memories, we analyzed the dynamics of synaptic weights within the population of neurons containing the overlapping memory sequences (i.e. neurons 200-249). Figure 6A, C, E shows distributions of synaptic weights for synapses pointing in the direction of S1 (top row) and in the direction of S1* (bottom row) before (blue) and after (red) specific events. Different columns correspond to different events, i.e. after S1 training (figure 6A, left), after S1* training (figure 6A, right), after sleep (figure 6C) or after interleaved training (figure 6E). Prior to any training, synaptic weights in the direction of either memory sequence were Gaussian distributed (figure 6A, blue histogram, left). After S1 training, the weights for S1 strengthened (shifted to the right), while the weights for S1* weakened (shifted to the left). As expected, this trend was reversed when S1* was trained (figure 6A, right). During sleep, for each sequence (S1 or S1*) there was a subset of synapses that were further strengthened, while the rest of synapses were weakened (figure 6C, red histogram). This suggests that sleep promotes competition between synapses, so specific subsets of synapses uniquely representing each memory trace can reach the upper bound to maximize recall performance while other synapses would become extinct to minimize interference. During interleaved training, we also observed separation of synapses but to a lesser extent than during sleep (figure 6E).

**Figure 6.**
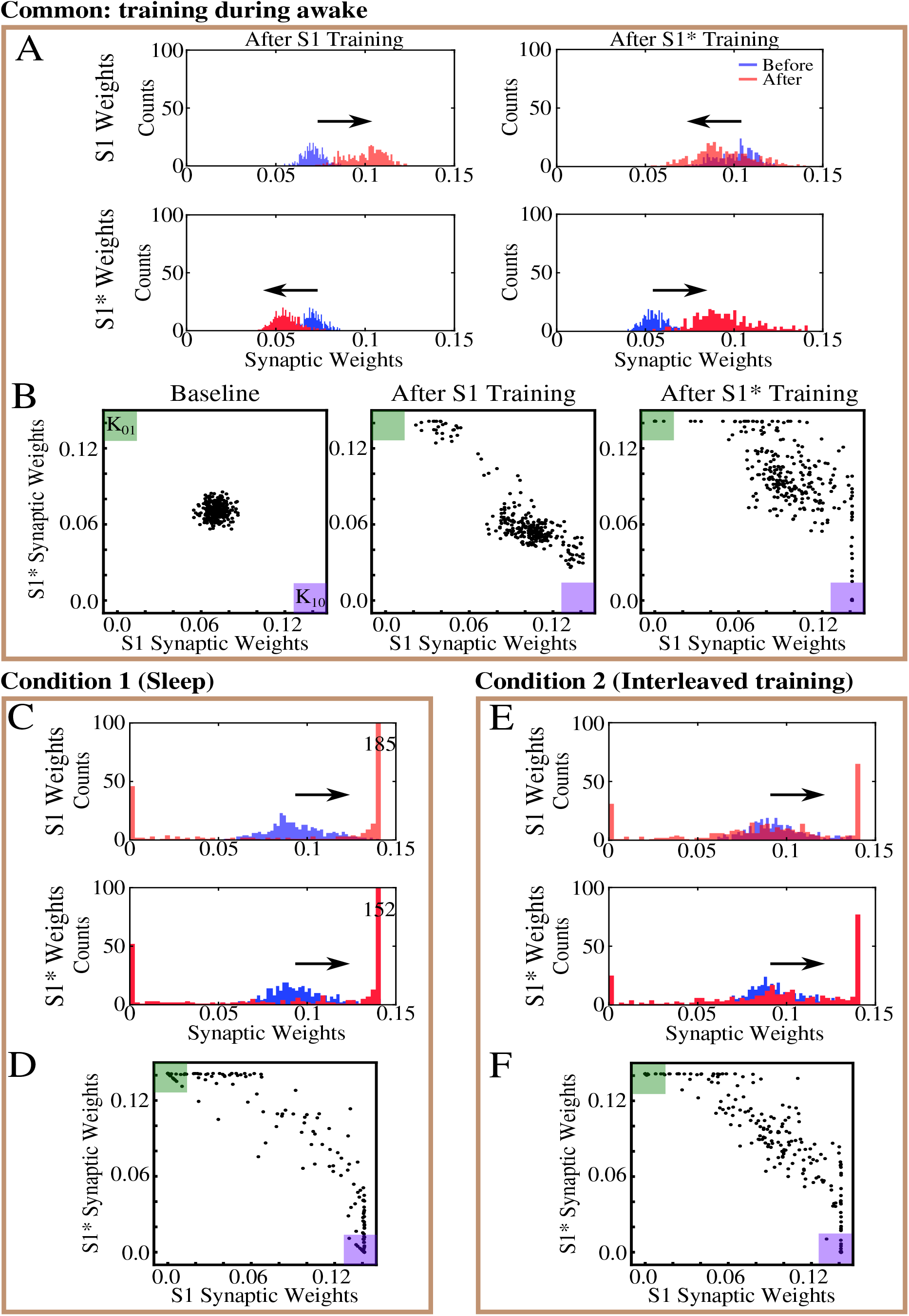
Sleep promotes unidirectional synaptic connectivity with different subsets of synapses becoming specific to the “old” and “new” memory sequences. **A**, Histogram of synaptic weights for all neuronal pairs (e.g., n1-n2) contributing to the memory sequences S1 and S1* after training of both memory sequences. Top row shows strength of synapses contributing to S1 only (e.g., n1->n2). Bottom row shows strength of synapses contributing to S1* only (e.g., n2->n1). Blue shows the starting points for weights, and red shows new weights. **B**, Scatter plots show synaptic weights for all pairs of neurons contributing to both S1 and S1* before and after training. For each pair of neurons (e.g., n1-n2), X-axis shows the strength of W_n1->n2_ synapse and Y-axis shows the strength of W_n2->n1_ synapse. The green (K_01_) and purple (K_10_) boxes represent the locations in the scatter plot where synaptic pairs which are strongly preferential for S1* (green) or S1 (purple) exist. **C/D**, Histograms of synaptic weights and scatter plots after sleep. **E/F**, Histograms of synaptic weights and scatter plots after interleaved training.

Because of the random anatomical connectivity, this cortical network model included two classes of synapses: *recurrent/bidirectional*, when a pair of neurons (e.g., n1 and n2) were connected by two synapses (n1->n2 *and* n2->n1) and *unidirectional* (n1->n2 *or* n2->n1). In the following we looked separately at these two classes to analyze the dynamics of synaptic weights during sleep vs interleaved training. In the scatter plots of S1*/S1 weights (figure 6B, D, F), for each pair of neurons (e.g., n1-n2), we plotted a point with the X-coordinate representing the weight of n1->n2 synapse and the Y-coordinate representing the weight of n2->n1 synapse. Therefore, any point with X- (Y-) coordinate close to zero would indicate a pair of neurons with functionally unidirectional coupling in S1* (S1) direction.

The initial Gaussian distribution of weights was pushed towards the bottom right corner of the plot (K_10_, purple box) indicating increases in S1 weights and relative decrease of S1* weights in response to S1 training (figure 6B, middle). Training of S1* caused an upward/left shift representing strengthening of S1* weights and weakening of S1 weights (figure 6B, right). For very long S1* training (not shown) almost all the weights would be pushed to the upper left corner (K_01_, green box). Sleep appears to have taken most of the weights located in the center of the plots (i.e., strongly bidirectional synapses) and separated them by pushing them to the opposite corners (K_01_, green box, and K_10_, purple box) (figure 6D). In doing so, sleep effectively converted strongly bidirectional connections into strongly unidirectional connections which preferentially contributed to the one memory sequence or another. During interleaved training, we also observed a trend for synaptic weights to move to the corners (i.e., to represent either one memory or another) but a much larger fraction of the weights remained in the middle of the diagram, and was even pushed towards (0,0) point indicating overall weakening of recurrent connections (figure 6F).

To further illustrate the dynamics of the recurrent synaptic weights, we projected the distributions of the weights from figure 6B, D, F to the “diagonal” K_01_-K_10_. We repeated this analysis for the model with interleaved training and compared the results with the sleep condition (figure 7). Before any training, the projection of synaptic weights exhibited a Gaussian distribution (figure 7A, left), reflecting the existence of functional recurrent synaptic connections in both sequence directions for most neuron pairs. Following S1 training, the distribution of the weights shifted in the direction of S1 indicating strengthening of synaptic weights in direction of S1 and a weakening of those in direction of S1* (figure 7A, middle). Next, S1* training strengthened recurrent weights in direction of S1* as indicated by the shift in the peak of distribution back towards the center (figure 7A, right). After a period of sleep, the distribution of synaptic weights became strongly bimodal. Almost complete lack of weights in the middle of distribution again suggests that most recurrent synaptic connections became functionally unidirectional thereby contributing to the storage of one sequence or the other but not both (figure 7C).

**Figure 7.**
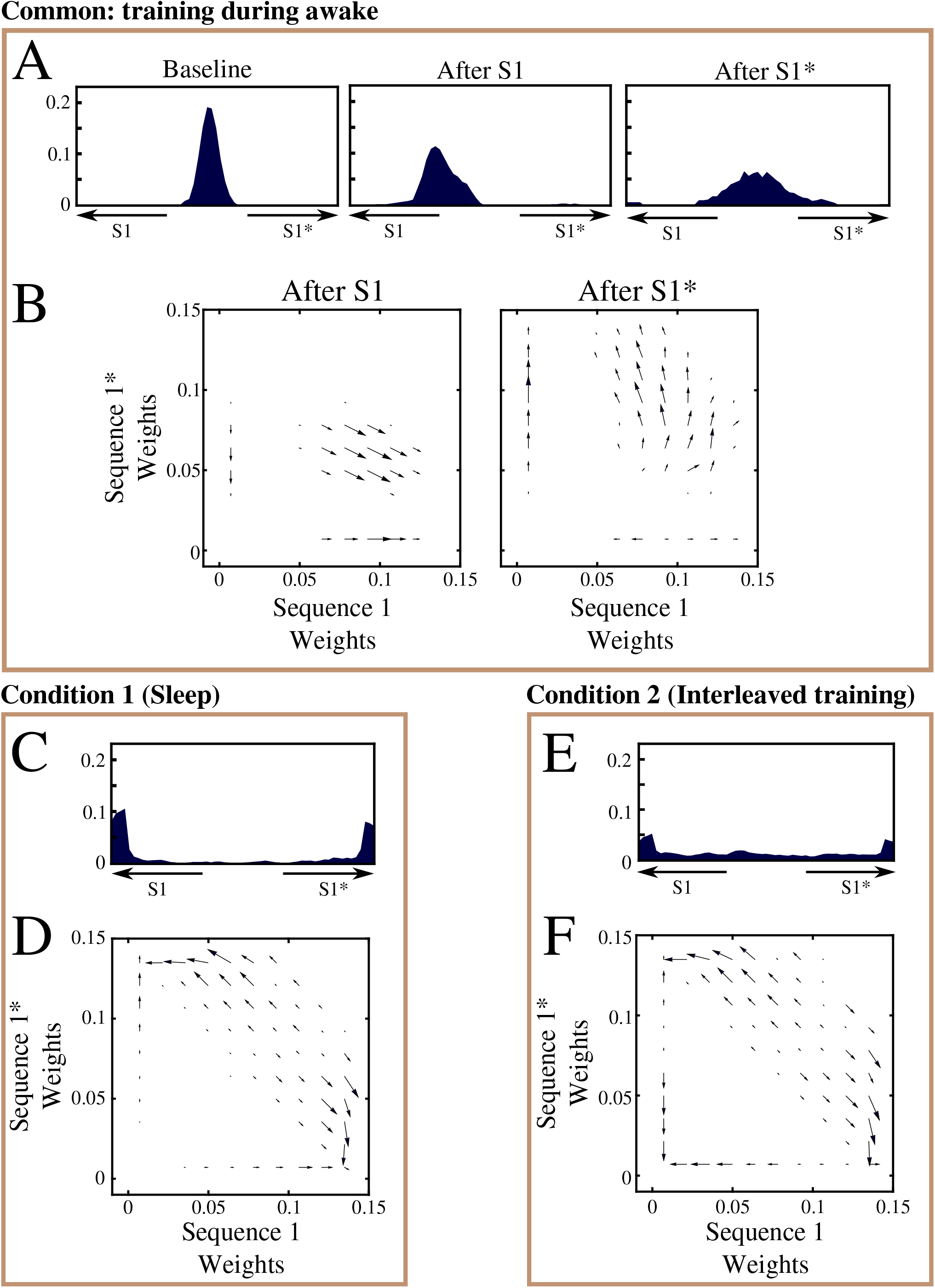
Both sleep and interleaved training lead to separation of synapses contributing to the “old” and “new” memory traces. **A/C/E**, Mean projection of synaptic weights for all pairs of neurons from the S1/S1* training region plotted in figure 6 B/D/E. X-axis – relative strength of S1 and S1* specific synapses for each pair of neurons. Middle of X-axis represents pairs with equal strength of S1 and S1* synapses. Extreme left (right) are pairs with functionally unidirectional synapses. Effects of either sleep or interleaved training are shown in C and E, respectively. Note the sharpening of the representation. **B/D/F**, Averaged synaptic dynamics showing evolution of synaptic weights during S1 and S1* training (B), sleep (D) and interleaved training (F).

The averaged dynamics of the synaptic weight changes are summarized in the vector plots (figure 7B, D, F). Note, these plots also include unidirectional connections before training – see vectors along the X- or Y-axis. The vector plots specific to the sleep phase revealed a separation of recurrent synaptic weights (figure 7D). Even relatively weak unidirectional synaptic weights were recruited for the storage of one sequence or another during sleep. This, however, was not the case for interleaved training. Figure 7E and F show that interleaved training also resulted in separation of recurrent synaptic weights. Unlike sleep, however, interleaved training preserved many more recurrent connections indicated by the thick band of weights seen in the middle of the weight projection (figure 7E). Increasing the duration of the interleaved training did not result in better separation of weights (data not shown). Interleaved training also suppressed weak unidirectional synaptic weights as shown in the vector plots (note arrows pointing to (0,0) in the bottom left area of the figure 7F. This was in stark contrast to the recruitment of the weak synaptic weights during sleep (compare to the figure 7D).

### Characteristic trends for the overall synaptic weights changes during sleep replay vs interleaved training

Figure 8 illustrates overall dynamics of synaptic weights in two models. During sleep, unidirectional connections increased in strength (note increase of mean strength and relatively small standard deviation in figure 8A1) while during interleaved training some unidirectional connections increased and others were suppressed (note no change of mean strength and large increase in standard deviation in figure 8A2). The mean strength of recurrent connections decreased slightly during both sleep and interleaved training (figure 8B), however the standard deviation increased much more during sleep (compare dashed lines in figure 8B1 vs B2). As discussed previously, the last could reflect strong asymmetry of the connections within recurrent pairs after sleep. To confirm this, we counted the total number of functionally recurrent and unidirectional connections after training and after sleep or after interleaved training (figure 8C). In this analysis if one branch of a recurrent pair reduced in strength below the threshold, it was counted as unidirectional. After sleep, the number of recurrent connections dropped to just about 15% of what it was after training (figure 8C1, green line), while after interleaved training it remained relatively high - about 50% of what it was initially (figure 8C2, green line). Decrease in the number of recurrent connections during sleep led to an increase in the number of functionally unidirectional connections (figure 8C, blue and red lines). Together these results indicate again that sleep decreases the density of recurrent connections and increases the density of unidirectional connections, both by increasing the *strength* of anatomical unidirectional connections and by *converting* anatomical recurrent connections to functionally unidirectional connections. This would minimize competition between synapses during recall of the overlapping S1 and S1* sequences. In contrast, interleaved training preserves but suppresses in strength a large fraction of the recurrent connections; it also cannot take full advantage of strengthening unidirectional connections. Below we show that this kind of dynamics leads to reduced transmission between populations of neurons representing different “letters” within a sequence and ultimately reduces performance.

**Figure 8.**
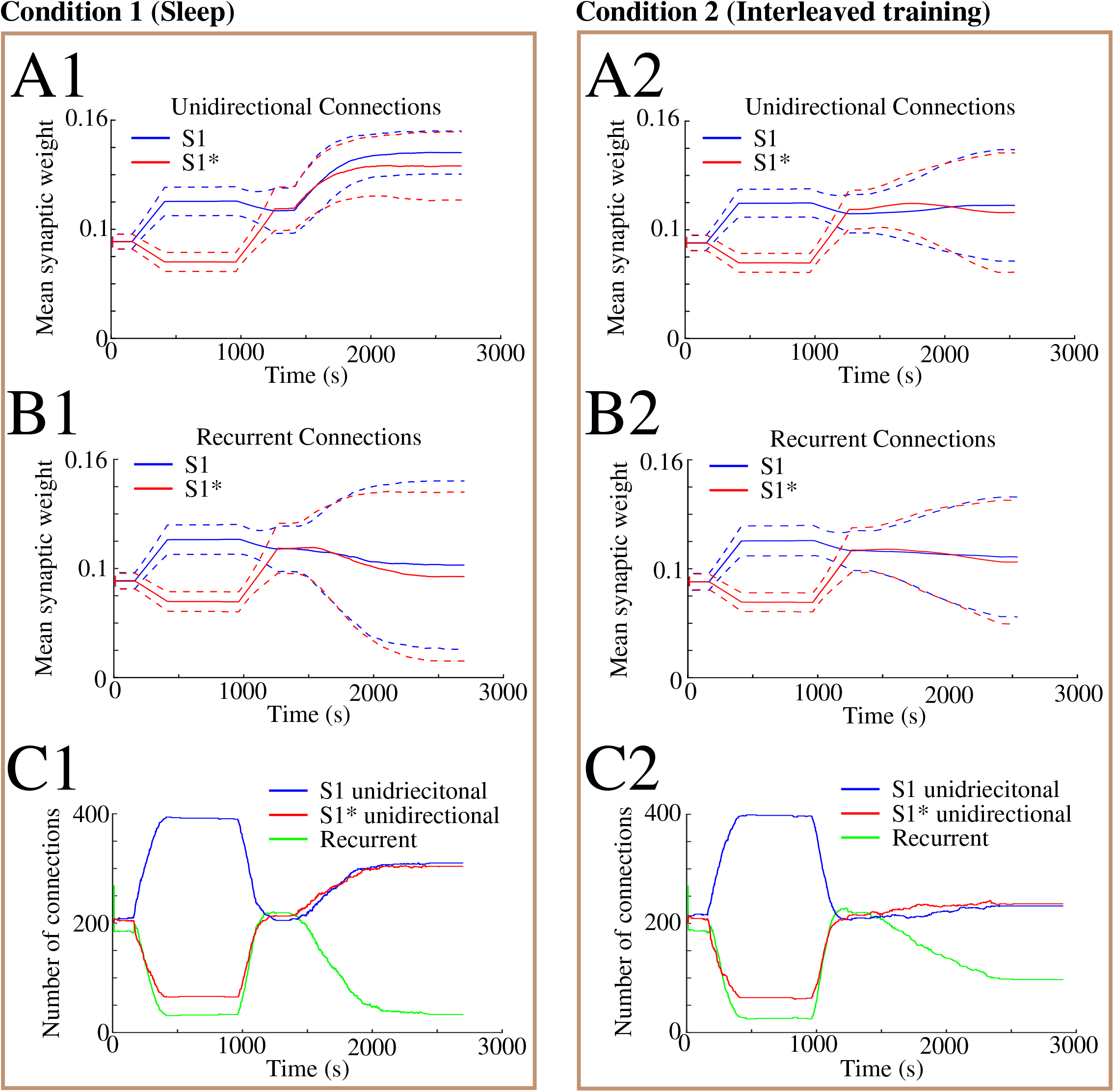
Overall synaptic weight changes during sleep vs interleaved training. In all panels index 1 corresponds to sleep condition, and index 2 corresponds to interleaved training. **A**, The evolution of the mean synaptic weight (solid line) and the standard deviation (dashed line) of unidirectional connections going along S1 (blue) and along S1* (red) in the trained region. Note overall increase in synaptic strength after sleep but not after interleaved training. **B**, The evolution of the mean synaptic strength (solid line) and the standard deviation (dashed line) of recurrent connections going along S1 (blue) and along S1* (red) in the trained region. Note large standard deviation after sleep indicating strong synaptic weight separation, so each recurrent neuronal pair supports either S1 or S1*. **C**, The number of recurrent and unidirectional connections in the trained region of the network as a function of time obtained after thresholding the network with threshold 0.065, which is smaller than the initial mean synaptic weight. Note decrease of functionally recurrent connections and increase of unidirectional connections after sleep.

### Sleep replay “fine tunes” synaptic connectivity to maximize separation of memory traces

To understand why different synaptic weight dynamics during sleep vs interleaved training lead to the difference in recall performance, we next evaluated the contribution of individual neurons to the recall of S1 vs S1* sequences. We assumed that a given neuron supports S1 if it receives strong input from the left (lower letter in sequence) and projects strongly to the right (higher letter in sequence); that is if a neuron n2 belongs to group B then it would support S1 if connections n1->n2 and n2->n3 are strong, for some n1 from group A and some n3 from group C. It would be opposite for neurons supporting S1*. For each neuron, we calculated the total “from left” + “to right” and “from right” + “to left” synaptic weights (see *Methods and Materials*) and plotted one against the other (figure 9A-C). On those plots, neurons placed closer to X (Y)-axis would be strongly supporting S1 (S1*) sequence and those in the middle, if they are sufficiently far from the origin, would support both sequences. As expected, initial training in awake had a tendency to create neurons supporting the most recently trained sequence. After training S1 the majority of neurons shifted to the right; S1* training reversed this trend (compare Fig 9 A left and right). After sufficiently long S1* training (not shown) the majority of neurons began supporting S1*. Both sleep and interleaved training had a tendency to “stretch” the neurons along the −45° diagonal (diagonal from coordinates (0,2) (2,0)) making both sequence *specific* and *generic* neurons (figures 9B, C), but sleep had larger fraction of “specialists” (neurons located close to X or Y axis) while interleaved training preserved more “generalist” neurons (located in the middle). Importantly, after interleaved training these neurons were moved on average closer to (0,0) point (weaker synaptic weights).

**Figure 9.**
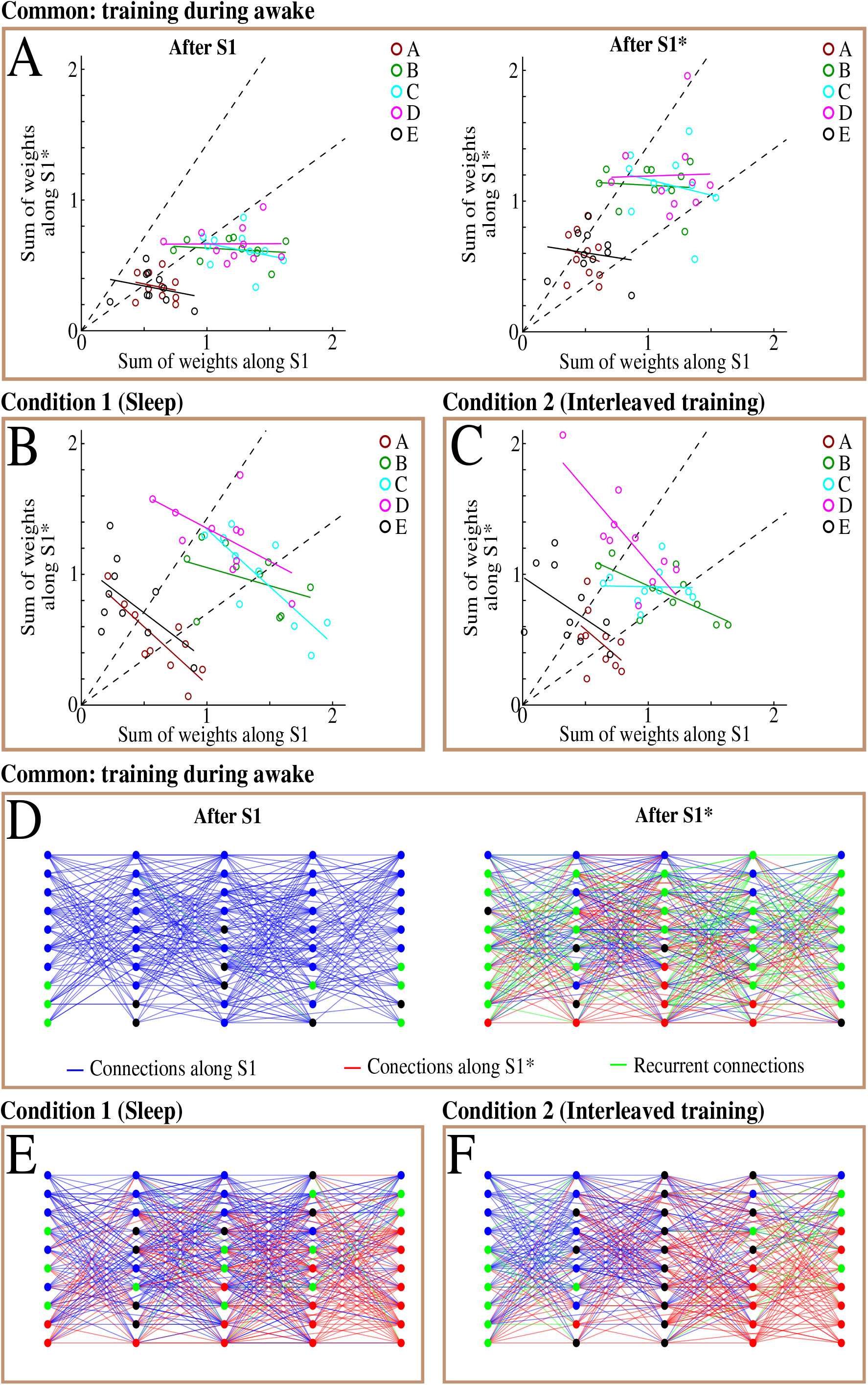
Sleep optimizes connectivity to reduce memory interference. The sum of weights of all incoming and outgoing connections to a neuron along S1 and S1* directions after training (**A**), after sleep (**B**), or, alternatively, after interleaved training (**C**). Each open circle corresponds to a neuron, and all neurons from the same group are colored alike; linear fit is obtained for each group separately and colored as neurons from the group. Dashed lines define boundaries of the regions, where neurons tend to represent a single sequence (bottom and top corners) or have a shared representation of both sequences (middle segment). **D**, Intergroup connectivity obtained after thresholding the network with threshold 0.08. Nodes are colored based on both total input and total output past a threshold for one sequence or the other (blue for S1 and red for S1*). If conditions are satisfied for both sequences, then the node is colored green. Otherwise, threshold conditions are failed for both directions, the node is colored black. Intergroup connectivity after sleep (**E**) and interleaved training (**F**).

The high sum of synaptic weights in one specific direction (figure 9A) may be driven by strong weights from one side and very weak weights from another (e.g., the sum of “from left” + “to right” synapses could be high because “from left” total synaptic weight is high but “to right” total synaptic weight is weak). To address this point, we looked at the evolution of the graph obtained by thresholding the weighted subgraph (see *Methods and Materials*) for neurons 200-249 from the region where interfering memory sequences were trained (figures 9D-F). To identify the role of each neuron, we colored neurons according to the following rule: a neuron was colored blue if *both* the total strength of “from left” and “to right” synapses was above the threshold defined by the strength value before any training. It was colored red if both the total strength of “from right” and “to left” synapses was above the threshold. It was colored green if both red and blue conditions were satisfied and it was colored black if none of the conditions were satisfied. The last would mean that the neuron was a “dead end” for both S1 and S1* sequences since at least one of two synaptic weights was below threshold for *both* directions.

Snapshots of the subgraph after S1 / S1* training and after sleep vs. interleaved training are shown in the figure 9D-F. S1 training first allocated all available connections for the purpose of S1 sequence completion (figure 9D, left panel, blue connections) and the majority of neurons became specialists for S1 (blue circles). The number of relatively strong recurrent synaptic connections increased after subsequent S1* training (figure 9D, right panel, green connections) and some connection became exclusively allocated to S1* (figure 9D, right panel, red connections). Many neurons changed their type to became generalists (green circles). After sufficiently long S1* training, all synaptic connectivity became reallocated to support S1* and the neurons became specialists for memory S1* (not shown), which would imply complete forgetting of S1. Sleep, presented after partial S1* training, enhanced both memories by decreasing recurrent connectivity in favor of either S1 (blue) or S1* (red) connections (figure 9E). Most of the neurons became specialists for one memory sequence or the other (red or blue circles) with some neurons remaining generalists (figure 9E). This trend was very different when interleaved training was applied, in place of sleep, as many connections remained recurrent (green lines; figure 9F). Furthermore, many neurons failed the test to support either one sequence or the other (see above) and were labeled black (especially in the middle group C). Those neurons had a reduced ability to support either one of the two memory sequences. This explains relatively lower memory completion performance after interleaved training (figure 4C) vs that after sleep (figure 5C).

It can be also noted that within each group, the neurons could be further separated into leaders and followers with respect to their response times (data is not shown). This suggests that once activity propagates from one group to another through a sequence specific set of inter-group connections, the rest of the group could be entrained through intra-group connectivity.

### Sleep forms memory specific synaptic connectivity profiles while preserving global communication between neurons

Previous observations led to our hypothesis that sleep attempts to reorganize network connectivity to an extreme configuration, where underlying connectivity is split into multiple separate groups: each representing only one memory sequence. To test this hypothesis, we first checked if the unweighted graphs obtained after thresholding of synaptic connectivity among neurons 200-249 had multiple maximal strongly connected components (MSCCs). Two neurons belong to the same MSCC if there is a directed path from one neuron to another along the directed edges of the unweighted graph. So, a single MSCC implies that there is a strong path between any two neurons in the network. We counted the evolution of the number of MSCCs for different threshold values on synaptic weights as shown in figure 10A. For a medium/high threshold (0.08), every neuron constituted an MSCC before training (figure 10A, orange) as the number of relatively strong connections was negligible because the mean of the initial synaptic weights was smaller than the threshold (see *Methods and Materials*). As training of S1 progressed, relatively strong connections were formed, which decreased the number of MSCCs. However, by the end of S1 training the number of MSCCs increased back to a value similar to that prior to training. This corresponded to an extreme reorganization of the connectivity in the trained region, where most of the relatively strong connections had a direction similar to the direction of S1. Therefore, the network became effectively disconnected in the opposite direction (hence low performance for S1*). After subsequent S1* training, there was a single MSCC again as there were many relatively strong connections in the network going in both directions, and this remained to be true after either sleep or interleaved training. If, however, S1* training continues in awake long enough, it would reallocate all connections to S1* and we would observe multiple MSCCs again (because the network is effectively disconnected in S1 direction) (not shown).

**Figure 10.**
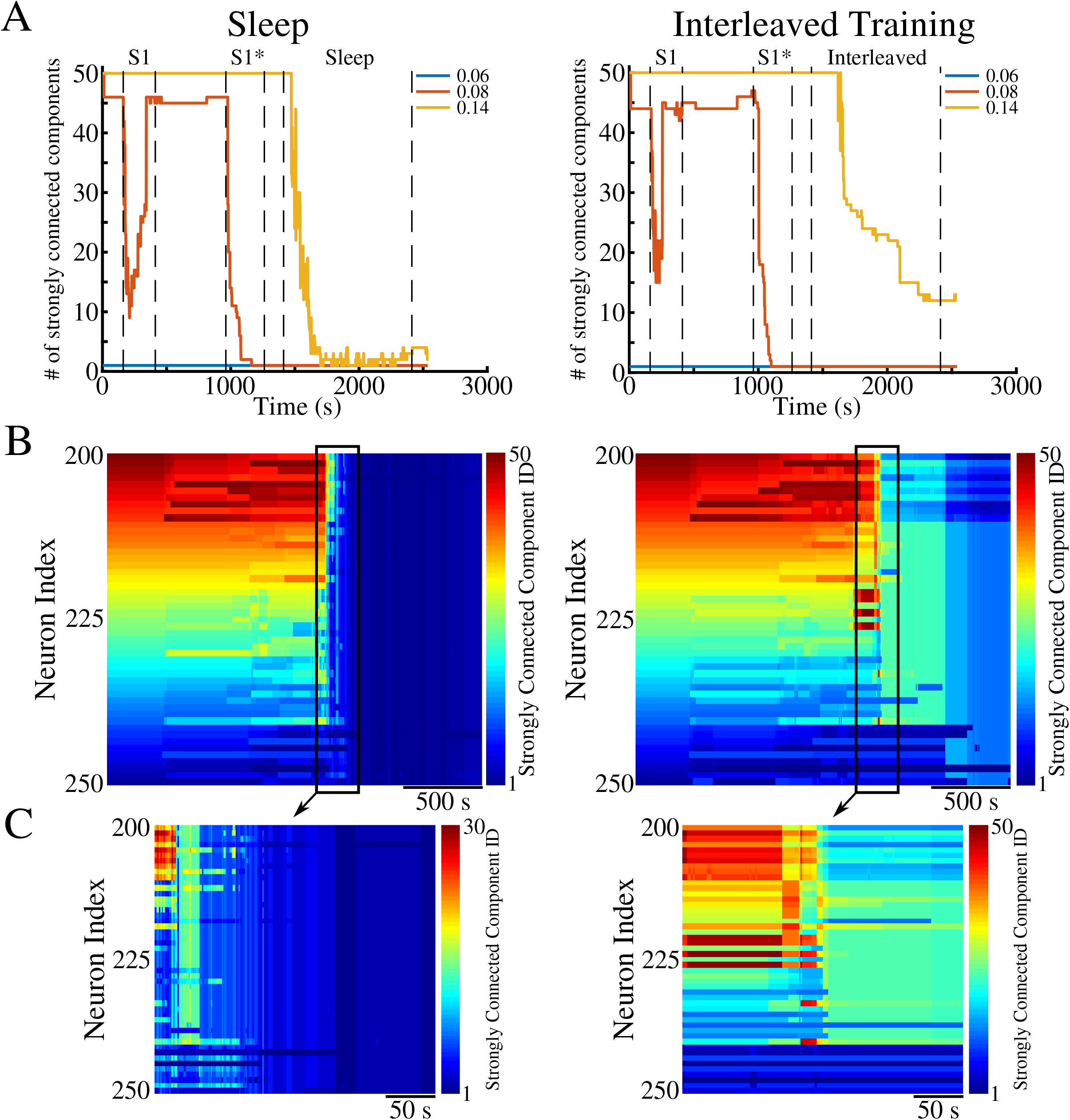
Reorganization of the cortical connectivity profile during sleep and interleaved training. Panels on the left correspond to sleep condition, while panels on the right correspond to interleaved training. **A**, The number of maximal strongly connected components (MSCC) in unweighted directed graphs obtained by thresholding the connectivity of the trained region in the cortex at every second of the simulation with thresholds 0.06 (blue), 0.08 (red), and 0.14 (orange). **B**, Neuronal MSCC membership in network color-coded in time through the entire duration of the simulations. **C**, Zoom-in of neuronal MSCC membership showing a drastic decrease in the number of MSCCs.

For a higher threshold value (0.14), which was equal to the upper bound on synaptic weights, the qualitative picture changed (figure 10A, yellow). At this level of granulation, the edges present in the unweighted graph after thresholding depict synapses that uniquely represented the trained sequences. There were not enough of these strong edges in the underlying graph after training of S1 or S1*, but they were created during sleep. While the unweighted graph had a small number of MSCCs for this threshold after sleep (figure 10A, left), the unweighted graph that resulted after interleaved training had a number of MSCCs exceeding the group size of a sequence, leading to segmentation of the connectivity profile into multiple MSCCs (figure 10A, right). Color-coded MSCC membership of each neuron from the trained region over the whole simulation duration is shown in figure 10B, and the region of significant connectivity reorganization is expanded in figure 10C. Figure 10B ruled out the possibility of a connectivity profile split into two subnetworks after sleep as MSCCs consisted of a giant MSCC, where most of the neurons belonged, and a small number of isolated neurons in remote parts of the trained region. That is, our model predicts that after sleep the cortical network reaches a state when memories are stored efficiently, i.e. without destroying communication between almost all neurons in the trained region, and at the same time, avoiding interference. The presence of a large number of relatively strong recurrent connections along with the segmentation of the connectivity profile formed by the strong edges into multiple MSCCs after interleaved training, and absence of these properties after sleep, may explain the difference in the performance of the two networks shown in figures 4C and 5C, respectively.

## Discussion

We report here that sleep can reverse catastrophic forgetting of previously trained (old) memories after damage by new training. Sleep was able to accomplish this task by spontaneous reactivation (replay) of both old and new memories without use of the original training data. The mechanism by which intrinsic network activity during sleep was able to recover and enhance memories was through reorganization and fine-tuning of the synaptic connectivity. In doing so, sleep created distinct synaptic weight representations of the overlapping memories that allow co-existence of the different memory traces without interference within overlapping ensembles of neurons. Thus, if without competition, a memory is represented by the entire available population of neurons, in the presence of competition, its representation is reduced to a *subset* of neurons who selectively and efficiently represents a given memory trace. This was achieved through memory replay during sleep, which (a) reduced reciprocal synaptic connectivity in favor of the functional unidirectional connections supporting one memory or another, and (b) increased strength of the anatomical unidirectional connections to better support individual memories. Our study predicts that after each new episode of training followed by sleep, representation of the old memories, sharing neurons with the new task, may change to achieve optimal separation of the old and new memory traces.

We compared sleep replay with interleaved training, that was previously proposed as a solution for catastrophic forgetting problem. Memory performance after interleaved training improved but was inferior to that after sleep replay. Our study suggests that sleep, by being able to directly reactivate memory traces encoded in synaptic weights patterns, provides a powerful mechanism to prevent catastrophic forgetting and enable continual learning.

### Catastrophic forgetting and continual learning

The work on catastrophic forgetting or interference in connectionist networks was pioneered by McCloskey and Cohen (Mccloskey and Cohen, 1989) and Ratcliff (Ratcliff, 1990). Catastrophic interference is observed when a previously trained network is required to learn new data, e.g., a new set of patterns. When learning new data, the network can suddenly erase the memory of the old, previously learned patterns (French, 1999; Hasselmo, 2017; Kirkpatrick et al., 2017). This type of forgetting of previously learned data occurs only after sufficient presentations of the new patterns. Catastrophic interference is thought to be related to the so-called “plasticity-stability” problem. The problem comes from the difficulty of creating a network with connections which are plastic enough to learn new data, while stable enough to prevent interference between old and new training sets. Due to the inherent trade-off between plasticity and memory stability, catastrophic interference and forgetting remains to be a difficult problem to overcome in connectionist networks (French, 1999; Hasselmo, 2017; Kirkpatrick et al., 2017).

A number of attempts have been made to overcome catastrophic interference (French, 1999; Hasselmo, 2017; Kirkpatrick et al., 2017). Early attempts included changes to the backpropagation algorithm, implementation of a “sharpening algorithm” in which a decrease in the overlap of internal representations was achieved by making hidden-layer representations sparse, or changes to the internal structure of the network (French, 1999; Hasselmo, 2017; Kirkpatrick et al., 2017). These attempts were able to reduce the severity of catastrophic interference in specific cases but could not provide a complete and generic solution to the problem. Another popular method for preventing interference or forgetting is to explicitly retrain or rehearse all the previously learned pattern sets while training the network on the new patterns – interleaved training (Hasselmo, 2017). This idea recently led to the number of successful algorithms to constrain the catastrophic forgetting problem, including generative algorithms to generate previously experienced stimuli during the next training period (Li and Hoiem, 2018; van de Ven and Tolias, 2018) and generative models of the hippocampus and cortex to generate examples from a distribution of previously learned tasks in order to retrain (replay) these tasks during a sleep phase (Kemker and Kanan, 2017).

In agreement with these previous studies, we show that interleaved training can prevent catastrophic forgetting resulted from training of the overlapping spike patterns sequentially. This method, however, does not change synaptic representations of the old memories to achieve optimal separation between old and new overlapping memory traces. Indeed, interleaved training requires repetitive activation of the entire memory patterns, so if different memory patterns compete for synaptic resources (as for the sequences studied here) each phase of interleaved training will enhance one memory trace but damage the others. This is in contrast to replay during sleep when only memory specific subsets of neurons are involved in each replay episode. On the synapse/cell level, we found that interleaved training preserved many recurrent synaptic pairs, however synaptic weights were reduced, leading to some fraction of “dead end” neurons which could not effectively contribute to memory recall. As a result, recall performance after interleaved training was saturated below that after sleep regardless of the training duration.

Another primary concern with interleaved training is that it becomes increasingly difficult/cumbersome to retrain all the memories as the number of stored memories continues to increase and access to the earlier training data may no longer be available. As previously mentioned, biological systems have evolved a mechanism to prevent this form of forgetting without the need to explicitly retrain the network on all previously encoded memories. Studying how these systems overcome this issue can provide insights into novel techniques to combat catastrophic forgetting in artificial neural networks.

### Sleep and memory consolidation

Though a variety of sleep functions remains to be understood, there is growing evidence for the role of sleep in consolidation of newly encoded memories (Oudiette and Paller, 2013; Paller and Voss, 2004; Rasch and Born, 2013; Stickgold, 2013; Walker and Stickgold, 2004; Wei et al., 2016; Wei et al., 2018). The mechanism by which memory consolidation is influenced by sleep is still largely debated, however a number of hypotheses have been put forward. One such hypothesis is the “Active System Consolidation Hypothesis” (Rasch and Born, 2013). Central to this hypothesis is the idea of repeated memory reactivation (Mednick et al., 2013a; Oudiette et al., 2013; Oudiette and Paller, 2013; Paller and Voss, 2004; Rasch and Born, 2013; Stickgold, 2013; Wei et al., 2016). Although NREM sleep was shown to be particularly important for consolidating declarative (hippocampus-dependent) memories (Marshall et al., 2006b; Mednick et al., 2013b), human studies suggest that NREM sleep may be involved in the consolidation of the procedural (hippocampus-independent) memories. This includes, e.g. simple motor tasks (Fogel and Smith, 2006), or finger-sequence tapping tasks (Laventure et al., 2016; Walker et al., 2002). Selective deprivation of N2 sleep, but not REM sleep, reduced memory improvement for the rotor pursuit task (Smith and MacNeill, 1994). Following a period of motor task learning, the duration of NREM sleep (Fogel and Smith, 2006) and the number of sleep spindles (Morin et al., 2008) increased. The amount of performance increase in the finger tapping task correlated with the amount of NREM sleep (Walker et al., 2002), spindle density (Nishida and Walker, 2007) and delta power (Tamaki et al., 2013). In a recent animal study (Ramanathan et al., 2015) consolidation of the procedural (skilled upper-limb) memory depended on the bursts of spindle activity and the waves of slow oscillation during NREM sleep.

### Model limitations and predictions

The model of training and sleep consolidation proposed in our new study has a close resemblance to learning and consolidation of procedural memory tasks, such that training a new task directly affects cortical synaptic connectivity that may be already allocated for other (older) memories. We found that as long as a damage to the older memory is not sufficient to completely erase its synaptic traces, sleep can enable replay of both older and newer memory traces and reverse the damage while improving performance. Thus, to avoid irreversible damage, new learning in our model is assumed to be slow which may correspond to learning a new motor skill over multiple days allowing sleep to recover older memory traces that are damaged by each new episode of learning.

We suggest that our model predictions, at least at the level of individual synapses and neurons, are not limited to specific type of memory (procedural) or specific type of sleep (NREM sleep). Indeed, we found that performance changes after sleep in the model were not dependent on the number of slow-waves but directly correlated with a total time in Up states. This suggests that replay during REM sleep (Louie and Wilson, 2001) may trigger synaptic weight dynamics similar to that we described here. While this model lacks hippocampal input, we showed previously (Sanda et al., 2019; Wei et al., 2016) that sharp wave-ripple like input to the cortical network would trigger replay of previously learned cortical sequences. This suggests, in agreement with the hippocampal indexing theory (Skelin et al., 2019), that replay driven by hippocampal inputs could reorganize the cortical synaptic connectivity in a matter similar to spontaneous replay we described here.

Our model of sleep consolidation predicts that the critical synaptic weights information from the previous learning is still preserved after new training even if recall performance for the older tasks is significantly reduced. Because of that, spontaneous activity during sleep can trigger reactivation of the previously learned memory patterns reversing damage. It further suggests that apparent loss of performance for older tasks in the artificial neuronal networks after new training – catastrophic forgetting - may not imply irreversible loss of information as it is generally assumed. Indeed, our recent work (Krishnan, 2019) revealed that simulating a sleep-like phase in artificial networks trained using backpropagation can provide a solution for catastrophic forgetting problem in agreement with our results from the biophysical model presented here. Few changes to the network properties, normally associated with transition to sleep, were critical to accomplish this goal: relative hyperpolarization of the pyramidal neurons and increasing strength of excitatory synaptic connectivity. Both are associated with known effects of neuromodulators during wake-sleep transitions (McCormick, 1992) and were implemented in the thalamocortical model (Krishnan et al., 2016) that we used in our new study. Interestingly, these changes would make neurons relatively less excitable and, at the same time, increase contribution of the strongest synapses, effective enhancing the dynamical range for the trained synaptic patterns and reducing contribution of synaptic noise; together this would promote replay of the previously learned memories during sleep.

To summarize, our model predicts that slow-wave sleep could prevent catastrophic forgetting and reverse memory damage through replay of the old and new memory traces. By selectively replaying new and competing old memories, sleep dynamics not only achieve consolidation of the new memories but also provides a mechanism for reorganizing synaptic connectivity responsible for the previously learned memories – re-consolidation of the old memory traces – to maximize separation between memories. By assigning different subsets of neurons to primarily represent different memory traces, sleep tends to orthogonalize memory representations to allow for the overlapping populations of neurons to store competing memories and to enable continual learning in the biological systems.

## Acknowledgements

This work was supported by the Lifelong Learning Machines program from DARPA/MTO (HR0011-18-2-0021) and ONR (MURI: N00014-16-1-2829).

## Author Contributions

O.C.G, Y.S., G.P.K, and M.B conceived the work. O.C.G, Y.S., and G.P.K performed the simulations and analysis. O.C.G, Y.S., G.P.K, and M.B wrote the manuscript.

## Methods and Materials

### Thalamocortical network model

#### Network architecture

The thalamocortical network model used in this study has been previously described in detail (Krishnan et al., 2016; Wei et al., 2016; Wei et al., 2018). Briefly, our network was comprised of a thalamus which contained 100 thalamocortical relay neurons (TC) and 100 reticular neurons (RE), and a cortex containing 500 pyramidal neurons (PY) and 100 inhibitory interneurons (IN). Our model contained only local network connectivity as described in figure 1. Excitatory synaptic connections were mediated by AMPA and NMDA connections, while inhibitory synapses were mediated through GABA_A_ and GABA_B_. Starting with the thalamus, TC neurons formed AMPA connections onto RE neurons with a connection radius of 8 (R_AMPA(TC-RE)_ = 8). RE neurons then projected inhibitory GABA_A_ and GABA_B_ connections onto TC neurons with R_GABA-A(RE-TC)_ = 8 and R_GABA-B(RE-TC)_ = 8. Inhibitory connections between RE-RE neurons were mediated by GABA_A_ connections with R_GABA-A(RE-RE)_ = 5. Within the cortex, PY neurons formed AMPA and NMDA connections onto PYs and INs with R_AMPA(PY-PY)_ = 20, R_NMDA(PY-PY)_ = 5, R_AMPA(PY-IN)_ = 1, and R_NMDA(PY-IN)_ = 1. PY-PY AMPA connections had a 60% connection probability, while all other connections were deterministic. Cortical inhibitory IN-PY connections were mediated by GABA_A_ with R_GABA-A(IN-PY)_ = 5. Finally, connections between thalamus and cortex were mediated by AMPA connections with R_AMPA(TC-PY)_ = 15, R_AMPA(TC-IN)_ = 3, R_AMPA(PY-TC)_ = 10, and R_AMPA(PY-RE)_ = 8.

#### Intrinsic currents

All neurons were modeled with Hodgkin-Huxley kinetics. Cortical PY and IN neurons contained dendritic and axo-somatic compartments as previously described (Wei et al., 2018). The membrane potential dynamics were modeled by the following equation:

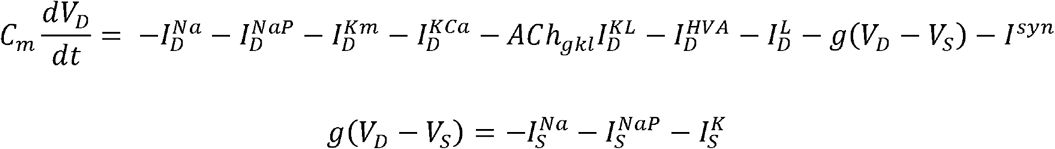

where *C*_*m*_ is the membrane capacitance, *V*_*D,S*_ are the dendritic and axo-somatic membrane voltages respectively, *I*^*Na*^ is the fast sodium (Na^+^) current, *I*^*NaP*^ is the persistent Na^+^ current, *I*^*Km*^ is the slow voltage-dependent non-inactivating potassium (K^+^) current, *I*^*KCa*^ is the slow calcium (Ca^2+^)-dependent K^+^ current, *ACh*_*gkl*_ represents the change in K^+^ leak current *I*^*KL*^ which is dependent on the level of acetylcholine (ACh) during the different stages of wake and sleep, *I*^*HVA*^ is the high-threshold Ca^2+^ current, *I*^*L*^ is the chloride (Cl^−^) leak current, *g* is the conductance between the dendritic and axo-somatic compartments, and *I*^*syn*^ is the total synaptic current input to the neuron. IN neurons contained all intrinsic currents present in PY with the exception of the *I*^*NaP*^.

Thalamic neurons (TC and RE) were modeled as single compartment neurons with membrane potential dynamics mediated by the following equation:

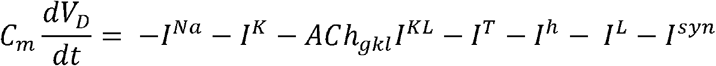

where *I*^*Na*^ is the fast Na^+^ current, *I*^*K*^ is the fast K^+^ current, *I*^*KL*^ is the K^+^ leak current, *I*^*T*^ is the low-threshold Ca^2+^ current, *I*^*h*^ is the hyperpolarization-activated mixed cation current, *I*^*L*^ is the Cl^−^ leak current, and *I*^*syn*^ is the total synaptic current input to the neurons. The *I*^*h*^ was only expressed in the TC neurons and not the RE neurons. The influence of histamine (HA) on *I*^*h*^ was implemented as a shift in the activation curve by *HA*_*gh*_ as described by:

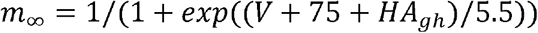

A detailed description of the individual currents can be found in our previous studies (Krishnan et al., 2016; Wei et al., 2018).

#### Synaptic currents and spike-timing dependent plasticity (STDP)

AMPA, NMDA, and GABA_A_ synaptic current equations were described in detail in our previous studies (Krishnan et al., 2016; Wei et al., 2018). The effects of ACh on GABA_A_ and AMPA synaptic currents in our model are described by the following equations:

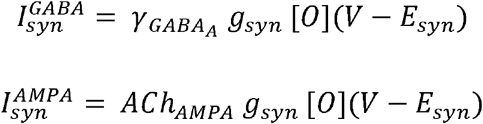

where *g*_*syn*_ is the maximal conductance at the synapse, [*O*] is the fraction of open channels, and *E*_*syn*_ is the channel reversal potential (E_GABA-A_ = −70 mV, E_AMPA_ = 0 mV, and E_NMDA_ = 0 mv). Parameter 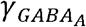 modulates the GABA synaptic currents for IN-PY, RE-RE, and RE-TC connections. For IN neurons 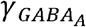 was 0.22 and 0.44 for awake and N3 sleep, respectively; 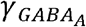 for RE was 0.6 and 1.2 for awake and N3 sleep. *ACh*_*AMPA*_ defines the influence of ACh levels on AMPA synaptic currents for PY-PY, TC-PY, and TC-IN. *ACh*_*AMPA*_ for PY was 0.133 and 0.4332 for awake and N3 sleep. *ACh*_*AMPA*_ for TC is 0.6 and1.2 for awake and N3 sleep.

Potentiation and depression of synaptic weights between PY neurons were regulated by spike-timing dependent plasticity (STDP). The changes in synaptic strength (g_AMPA_) and the amplitude of miniature EPSPs (A_mEPSP_) have been described previously (Wei et al., 2018):

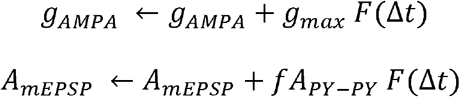

where *g*_*max*_ is the maximal conductance of *g*_*AMPA*_, and f = 0.01 represents the slower change of STDP on *A*_*mEPsP*_ as compared to *g*_*AMPA*_. The STDP function F is dependent on the relative timing (Δt) of the pre- and post-synaptic spikes and is defined by:

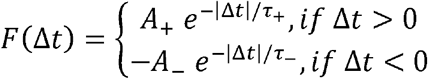

where A_+/−_ set the maximum amplitude of synaptic change. A_+/−_ = 0.002 and τ_+/−_ = 20 ms. A_−_ was reduced to 0.001 during training to reflect the effects of changes in acetylcholine concentration during focused attention on synaptic depression during task learning observed experimentally (Blokland, 1995; Shinoe et al., 2005; Sugisaki et al., 2016).

#### Sequence training and testing

Training and testing of memory sequences was performed similar to our previous study (Wei et al., 2018). Briefly, trained sequences were comprised of 5 groups of 10 sequential PY neurons. Each group of 10 were sequentially activated by a 10 ms DC pulse with 5 ms delay between subsequent group pulses. This activation scheme was applied every 1 s throughout the duration of the training period. Sequence 1 (S1) consisted of PY groups (in order of activation): A(200-209), B(210-219), C(220-229), D(230-239), E(240-249). Sequence 2 (S2) consisted of PY groups (in order of activation): W(360-369), V(350-359), X(370-379), Y(380-389), Z(390-399) and can be referred as non-linear due to the non-spatially sequential activations of group W, V, and X. Sequence 1* (S1*) was trained over the same population of neurons trained on S1 but in the reverse activation order (i.e. E-D-C-B-A). During testing, the network was presented with only the activation of the first group of a given sequence (A for S1, W for S2, and E for S1*). Performance was measured based on the network’s ability to recall/complete the remainder of the sequence (i.e. A-B-C-D-E for S1) within a 350 ms time window. Similar to training, test activation pulses were applied every 1 s throughout the testing period. Training and testing of the sequences occurred sequentially as opposed to simultaneously as in our previous study (Wei et al., 2018).

### Data analysis

All analyses were performed with custom built Matlab and Python scripts. Data are presented as mean ± standard error of the mean (SEM) unless otherwise stated. For each experiment a total of 10 simulations with different random seeds were used for statistical analysis.

#### Sequence performance measure

A detailed description of the performance measure used during testing can be found in (Wei et al., 2018). Briefly, the performance of the network on recalling a given sequence following activation of the first group of that sequence (see Methods and Materials: *Sequence training and testing*) was measured by the percent of successful sequence recalls. We first detected all spikes within the predefined 350 ms time window for all 5 groups of neurons in a sequence. The firing rate of each group was then smoothed by convolving the average instantaneous firing rate of the group’s 10 neurons with a Gaussian kernel with window size of 50 ms. We then sorted the peaks of the smoothed firing rates during the 350 ms window to determine the ordering of group activations. Next, we applied a string match (SM) method to determine the similarity between the detected sequences and an ideal sequence (ie. A-B-C-D-E for S1). SM was calculated using the following equation:

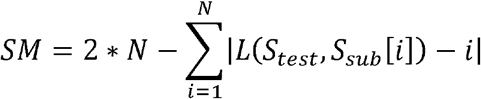

where N is the sequence length of S_test_, S_test_ is the test sequence generated by the network during testing, S_sub_ is a subset of the ideal sequence that only contains the same elements of S_test_, and *L*(*S*_*test*_,*S*_*sub*_[*i*]) is the location of the element S_sub_[i] in sequence S_test_. SM was then normalized by double the length of the ideal sequence. Finally, the performance was calculated as the percent of recalled sequences with SM≥Th, where Th is the selected threshold (here, Th = 0.8) indicating that the recalled sequence must be at least 80% similar to the ideal sequence to be counted as a successful recall as previously done in (Wei et al., 2018).

#### Sequence replay during N3 sleep

To find out whether a trained sequence is replayed in the trained region of the network during the Up state of a slow-wave in N3 sleep, we first identified the beginning and the end of each Up state by considering sorted spike times of neurons in each group. For each group, the time instances of consecutive spikes that occur within a 15 ms window were considered as candidate members of an Up state, where the window size was determined to decrease the chance of two spikes of the same neuron within the window. To eliminate spontaneous spiking activity of a group that satisfies the above condition but is not part of an Up state, we additionally required that the period between two upstate was at least 300 ms, which corresponds to a cortical Down state. The values for window durations reported above were identified to maximize the performance of the Up state search algorithm.

Once all Up states were determined, we defined the time instances when groups were active in each Up state. A group was defined as active if the number of neurons from the group that spikes during 15 ms exceeded the activation threshold, and the instance when the group is active was defined as the average over spike times of a subgroup of neurons with the size equals to the activation threshold within the 15 ms window. In our study the activation threshold was selected to be half of a group size (i.e. 5 neurons). Using sorted time instances when groups are active, we counted the number of times a possible transition between arbitrary groups, and if all four transitions of a sequence were observed sequentially in the right order then we counted that as a replay of the sequence.

#### Connectivity thresholding

To track connections in the trained region of a network, we performed thresholding of the underlying network, called weighted graph, at each second of simulation time. Thresholding results in a directed unweighted graph, where directed edges define significant edges of the weighted graph for storing the memories. The initial values of synaptic weights are drawn from the same Gaussian distribution with mean 0.0707 and variance 10% of the mean. A synaptic connection that encodes a sequence should undergo potentiation, and, hence, its strength should increase in such an event. With this in mind, we selected a threshold to be 0.08 in figure 9D, E, and F and figure 10 that would guarantee that a specific synapse has taken part in storing a memory. Using this relatively high threshold we looked at the connectivity profile of a directed unweighted graph. We dropped the directions of connections in figure 9D, E, and F, and instead, colored connections depending on whether they are unidirectional and go in one or the opposite direction, or there is a recurrent connection going in the opposite direction. To avoid a runaway of synaptic strength over entire simulation protocols, an upper bound on synaptic strength was imposed, which was selected to be twice the mean of initial weight. This upper bound was also used as a threshold. The directed unweighted graph obtained with this threshold value should identify connections which strongly support a memory, and this value was used in figure 10A-C.

#### Total synaptic weight analysis

For every neuron from a group we computed the total synaptic weight “from left” and “to right”, by considering the sum of all weights of synapses projecting on the neuron from neurons in preceding group, with respect to propagation of activity within a memory sequence if such a group exists, and the sum of all weights of synaptic connections from the neuron to the following group, if there is such a group. We omitted all synaptic connections within the group to which the neuron, for which the total synaptic weight is computed, belongs.

